# Change in social rank and brain dopamine levels: findings from a novel pig model

**DOI:** 10.1101/2020.08.06.239780

**Authors:** Hui G. Cheng, Abiy M. Mohammed, Anthony P. Pease, Joshua Gehrke, George Bohart, Robert Burnett, Michael A. Nader, James C. Anthony

**Author notes:** These two authors contributed equally (Hui G. Cheng, Abiy M. Mohammed). **Address correspondence to**, Hui G. Cheng, Ph.D., Department of Epidemiology and Biostatistics, Michigan State University, East Lansing, MI 48823.

## Abstract

Dopamine (DA) signaling is central in hypothesized causal paths linking the influence of social and environmental variables with cognition, behavior and affective states, including vulnerability to drug dependence. Here, we study whether *change in one’s social rank* induces DA and norepinephrine (NE) changes using a novel pig model with a social-ranking-and-re-ranking protocol to investigate social context influences on catecholamine concentrations in cerebrospinal fluid (CSF) and blood. For two weeks, 16 recently weaned male piglets were socially housed in four groups, with video-recordings for social rank assessments (α, β, γ, and δ); CSF and blood were obtained from these stable social groups. Next, all four α were housed together, as were all four β, etc., again with video recording for blinded social ranking. CSF and blood samples were collected at three time points: prior to initial social housing, following social housing and following re-organization. Regression analyses disclosed a positive relationship between changes in social rank and post-rank change in CSF levels of DA; one unit increase of social rank predicted a 17.4 pg/ml increase in CSF dopamine concentrations (95% CI= 1.2, 33.7). Compared to piglets with downward shifts in ranks (i.e., high-to-low), piglets with upward shifts (i.e., low-to-high) had a statistically significant greater increase in CSF DA levels. No relationship was observed for CSF NE or blood concentrations of DA or NE at any phase of this experiment. This work, using a novel pig model, adds new evidence on alteration of the brain dopaminergic system induced by social rank change.

## 1. Introduction

Social status is a central facet in humans, one of the few consistent and strong predictors of health and disease, such that it has been named a ‘fundamental cause of diseases’ (Link and Phelan, 1995; Marmot, 2005). In humans, living in economically disadvantaged communities, being bullied, and abrupt downward shifts in social rank (e.g., becoming unemployed and economic misfortune) have well-documented associations with the occurrence of dopamine-mediated disturbance, including alcohol and drug dependence (Dohrenwend et al., 1992; Hemmingsson et al., 1999; Nobile et al., 2007). Nonetheless, human studies have a limited capability to disclose hypothesized molecular mechanisms for the effects of social rank on the course of alcohol and drug dependence due to ethical considerations in humans.

Studies in nonhuman primates have provided important evidence on social rank altering the central nervous system at a molecular level and predisposing individuals with lower social rank to higher risks of alcohol and drug problems compared to those with higher social rank (Crowley et al., 1992, 1974; Czoty et al., 2005; Morgan et al., 2002). Using positron emission tomography to assess pre-social housing dopamine (DA) D2/D3 receptor availability (sometimes referred to as binding potential), a series of studies have suggested that social rank influences the rates of cocaine self-administration, across a wide range of doses, via alterations of DA D2/D3 receptor binding potential. When initially tested, cocaine served as a reinforcer in subordinate monkeys, but not in dominant monkeys (Morgan et al., 2002), and following extensive cocaine exposure, when the conditions were changed to a cocaine-food choice paradigm, subordinate monkeys were still more sensitive to cocaine than dominant monkeys (Czoty et al., 2005). An implicated underlying mechanism involves the acquisition of a higher social rank serving the same function as an alternative reinforcer to cocaine and altering DA D2/D3 receptor availability and perhaps activity of endogenous DA such that exogenously administered cocaine did not initially function as a reinforcer (Morgan et al., 2002) and subsequently attenuating cocaine reinforcement (Nader and Woolverton, 1991).

The initial differences in D2/D3 receptor binding potentials between dominant and subordinate monkeys were no longer apparent after extensive cocaine self-administration histories (Czoty et al., 2004).

Importantly for the present study, during abstinence from cocaine, differences in D2/D3 receptor binding potentials re-emerged, with dominant animals having significantly higher receptor availability compared to subordinate monkeys (Czoty et al., 2004), suggesting that D2/D3 receptor availability is malleable with orderly changes observable following environmental and pharmacological manipulations. Similarly, in female monkeys, D2/D3 receptor availability changed in dominant monkeys when placed in social groups and returned to baseline when monkeys were again individually housed (Michael A. Nader et al., 2012).

Social rank in humans is often in a dynamic change throughout one’s lifetime (e.g., ascents or drops in popularity, academic or work performance, athletic competence, etc.). However, it remains to be determined whether changes in social status (rank) can result in changes in the brain DA system in humans and influence DA-mediated behaviors, such as drug abuse. Moreover, whereas previous nonhuman primate studies have shown that D2/D3 receptor availability is not different in monkeys prior to social housing (i.e., not a trait marker; Morgan *et al*, 2002; Nader *et al*, 2012b), other unobserved heterogeneities may account for the relationship between social rank and the dopaminergic system (e.g., early environment, aggression, novelty-seeking, etc.; Morgan *et al*, 2000; Tung *et al*, 2011). More definitive evidence can be secured in a crossover ranking-and-re-ranking research design. In the crossover design, the focal point is the change in social rank within each individual, and therefore minimizes between-subject variations. Whether changes in social rank can influence DA neurotransmission in drug-naïve individuals has not yet been determined and may be a better model of social fluctuations that occur prior to initial drug exposure that could influence vulnerability to substance abuse.

In this report, we describe a novel crossover ranking-and-re-ranking experiment in domestic pigs (*Sus Scrofa*), a potentially suitable species with a recently annotated genome, DA system and other brain analogies to nonhuman and human primates, and existing drug self-administration lines of research (Gieling et al., 2011). The central hypothesis is that post-ranking brain DA level will increase with each unit increase of social rank at re-ranking.

## 2. Results

### Initial Social Rank Determinations

Pre-social housing CSF concentrations are shown in Table 1. Baseline weight or catecholamine concentrations in blood or CSF did not predict the first social rank (regression coefficient = −0.002 to 0.195; all p-value > 0.09; Table 2). In addition, body weight was not associated with CSF or plasma measures of DA and NE at baseline (correlation coefficient ρ=−0.50 for CSF DA, p=0.085; ρ= − 0.28 for CSF NE, p=0.362; ρ=0.39 for plasma NE, p=0.139; ρ= −0.18 for plasma epinephrine, p=0.509; ρ=0.03 for plasma DA, p=0.928). After stable social groups were established (Day 14), the initial social rank was associated with smaller increases in CSF and blood NE concentrations (CSF NE: regression coefficient=− 125.7pg/ml, 95% CI= −241.1, −10.3, p=0.039, n=9; blood NE: regression coefficient=−139.8pg/ml, 95% CI= − 276.9, −2.7, p=0.046, n=14). The initial social rank was not associated with changes in either CSF or blood DA concentrations (CSF DA: regression coefficient=24.0 pg/ml, 95% CI= −11.0, 59.0, p=0.149, n=9; blood DA: regression coefficient=6.0pg/ml, 95% CI= −19.2, 31.2, p=0.613, n=14).

**Table 1.** Results of linear regression (β) for baseline weight and catecholamine levels predicting the first social rank in pigs.

| Molecule | n | Baseline level | $\beta$ | 95% CI | p |
| --- | --- | --- | --- | --- | --- |
|  |  | mean (s.d.) |  |  |  |
| Weight (lb) | 16 | 14.7 (0.7) | 0.195 | -0.722, 1.112 | 0.656 |
| CSF DA (pg/ml) | 13 | 48.9 (39.6) | -0.010 | -0.028, 0.007 | 0.217 |
| CSF NE (pg/ml) | 13 | 275.6 (120.5) | 0.005 | -0.001, 0.010 | 0.093 |
| Blood DA (pg/ml) | 16 | 101.1 (39.7) | <0.001 | -0.001, <0.001 | 0.492 |
| Blood NE (pg/ml) | 16 | 923.9 (1026.6) | -0.002 | -0.016, 0.012 | 0.731 |
| Blood EPI (pg/ml) | 16 | 71.6 (46.7) | 0.001 | -0.016, 0.018 | 0.909 |

n, numbers of cases for which sample collection was succesful at baseline. DA, dopamine; NE, norepinephrine; EPI, epinephrine; s.d., standard deviation.

**Table 2.** Description of post-ranking changes in catecholamines and normality test.

| **Molecule** | **n** | **Mean change**<br>**(s.d.)** | **Skewness\*** | **Kurtosis\*** | **Joint test\*** |
| --- | --- | --- | --- | --- | --- |
| CSF DA (pg/ml) | 8 | 48.5 (38.3) | 0.736 | 0.864 | 0.931 |
| CSF NE (pg/ml) | 8 | 51.4 (199.0) | 0.410 | 0.494 | 0.524 |
| Blood DA (pg/ml) | 14 | 68.7 (58.3) | 0.468 | 0.863 | 0.746 |
| Blood NE (pg/ml) | 14 | 113.0 (190.4) | 0.880 | 0.053 | 0.130 |
| Blood EPI (pg/ml) | 14 | -5.1 (16.8) | 0.343 | 0.945 | 0.607 |

n, numbers of cases for which sample collection was succesful at baseline. DA, dopamine; NE, norepinephrine; EPI, epinephrine; s.d., standard deviation.
\* p values for the test for skewness, kurtosis, and a joint chi-square test.

### Social Reorganization

After the two rounds of ranking, four piglets stayed at the same social rank (e.g., the α that stays α in the all-α group and the β that stays β in the all-β group), six piglets had an increase in their social ranks (i.e., three with an one unit increase: δ to γ, γ to β, and β to α; two with a two-unit increase: δ to β, and γ to α; one with a three-unit increase: δ to α), and six piglets had decreased social ranks.

According to the joint test for skewness and kurtosis, no evidence was found for a non-normal distribution for changes in CSF and blood catecholamine levels as well as changes in social rank (p-values= 0.13 to 0.93 for various catecholamines and p-value=0.96 for changes in social rank, see Table 2).

According to linear regression estimates, there was a statistically significant positive relationship between changes in post-ranking CSF DA concentrations with changes in social rank (Table 3 and Fig. 2, Panel 1; β=17.4 pg/ml, 95% CI=1.2, 33.7). That is, for every one-unit increase in social rank, there was a mean increase of 17.4 pg/ml in the CSF DA concentrations. In contrast, no such relationship was found for CSF NE (Fig. 2, Panel 2) or blood DA or NE levels (Fig. 2, Panels 3 and 4; β=10.2 pg/ml, 95% CI= −113.4, 133.7 for CSF NE; β=−7.1 pg/ml, 95% CI= −31.3, 17.2 for blood DA, see Table 3). Statistical inference remains the same when including the pig with extreme blood catecholamine measures.

**Table 3.** Estimated regression coefficients (β) linking changes in the levels of catecholamines in cerebrospinal fluid (CSF) and blood with one unit change in social ranks following social re-ranking.

| **Molecules** | **n** | **$\beta$** | **95% CI** |
| --- | --- | --- | --- |
| CSF DA (pg/ml) | 8 | **17.4** | **1.2, 33.7** |
| CSF NE (pg/ml) | 8 | 10.2 | -113.4, 133.7 |
| Blood DA (pg/ml) | 14 | -7.1 | -31.3, 17.2 |
| Blood NE (pg/ml) | 14 | -50.3 | -124.2, 23.6 |
| Blood EPI (pg/ml) | 14 | -1.2 | -8.5, 6.2 |

n, numbers of cases for which sample collection was successful in both occasions. DA, dopamine; NE, norepinephrine; EPI, epinephrine.
Bold font indicates statistical significance at 0.05 level.

Exploratory analysis on differences in post-rank changes in catecholamine levels across high-and-low-rank groups revealed potential moderation by initial social rank (Table 4). Compared to piglets with downward shifts in ranks (i.e., high-to-low), piglets with upward shifts (i.e., low-to-high) had a statistically significant greater boost in CSF DA levels. No other comparisons showed statistically significant changes in CSF or blood catecholamines when compared to piglets with upward shifts in social rank.

**Table 4.** Estimated regression coefficients (β) linking changes in the levels of catecholamines in cerebrospinal fluid (CSF) and blood with change in social ranks stratified by initial social rank (reference= low-to-high).

| Molecules | n | low-to-low |  | high-to-low |  | high-to-high |  |
| --- | --- | --- | --- | --- | --- | --- | --- |
| | | $\beta$ | 95% CI | $\beta$ | 95% CI | $\beta$ | 95% CI |
| CSF DA (pg/ml) | 8 | -52.0 | -119.4,15.4 | **-67.5** | **-134.9,-0.1** | -69.0 | -154.3,16.3 |
| CSF NE (pg/ml) | 8 | -162.5 | -765.7,440.7 | 50.0 | -553.2,653.2 | -20.0 | -783.1,743.1 |
| Blood DA (pg/ml) | 14 | -57.3 | -135.3,20.8 | 42.5 | -41.8,126.8 | 20.8 | -63.5,105.1 |
| Blood NE (pg/ml) | 14 | 29.3 | -277.7,336.2 | 217.0 | -114.5,548.5 | 56.7 | -274.9,388.2 |
| Blood EPI (pg/ml) | 14 | -18.5 | -39.4,2.4 | 4.5 | -21.1,30.1 | 12.7 | -9.9,35.3 |

n, numbers of cases for which sample collection was successful in both occasions. DA, dopamine; NE, norepinephrine; EPI, epinephrine.
Bold font indicates statistical significance at 0.05 level.

## 3. Discussion

In this study, we report findings from an experiment in domestic pigs using a novel social-ranking-and-re-ranking design. Piglets were randomly selected from different litters which should control for any effect of pre-experiment ranking; neither pre-social-housing catecholamine concentrations in CSF or blood nor body weight predicted their initial social rank. Following social reorganization, we found evidence that changes in social rank induced changes in the brain dopaminergic system in a relatively short timeframe (Greene and Faull, 1989). More specifically, increases in social rank were associated with larger increases in CSF levels of DA. In contrast, no changes were observed for NE in CSF and blood or in blood DA levels as a result of changes in social rank, highlighting the specificity of social rank to the brain dopaminergic system.

Previous studies have shown that enriched or deprived environments can modulate developmental trajectories of DA levels, alcohol consumption, as well as aggressive and other socially maladaptive behaviors in nonhuman primates and rats (Bowling et al., 1993; Clarke et al., 1996; Harlow and Zimmermann, 1959; J D Higley et al., 1991). Findings from the present study extend previous research on social environment and DA-mediated changes in behavior and shed new light on the underlying mechanism for how social environment can modify an individual’s vulnerability to drug problems. An ascendance in social rank creates an enriched environment for the individual and can serve as a reinforcer; in contract, a decline in social rank creates a deprived environment and can be a stressor (Czoty *et al*, 2004; Koob and Le Moal, 1997; Volkow *et al*, 1999; Nader *et al*, 2012a). Both of these processes can induce changes to the dopaminergic system and play central roles in drug-use-related behaviors and other behavioral outcomes (Koob and Volkow, 2010; M A Nader et al., 2012; Volkow and Morales, 2015). In addition, according to our exploratory analysis, the largest effects are found for those who had an upward shift in social rank (i.e., low-to-high). This result is in line with findings from a previous study in nonhuman primates showing increased D2/D3 receptor availability in the caudate nucleus and putamen among individuals with initial low ranks before regrouping, compared to previously dominant monkeys (Czoty *et al*, under review). Taken together, these results suggest that the dopaminergic system is responsive to rewarding experiences in a relatively short time frame.

In this study, CSF DA levels increased for all but one piglet after the second round of ranking. Possible reasons for these increases include changes in the DA system associated with the development of piglets and changes induced by social interaction, in general, independent of social rank (Kanitz et al., 2009, 2004). These increases in DA are also consistent with a PET imaging study in individually housed adolescent monkeys showing age-dependent changes in D2/D3 receptor measures (Gill et al., 2012). Future studies using a similar experimental design in adult pigs will be able to elucidate the mechanisms mediating these DA changes. Nonetheless, we found greater increase in extracellular DA availability among piglets with increased social rank compared to those with decreased social rank. Such a finding is in line with work involving socially housed male and female cynomolgus monkeys (Morgan et al., 2002; Riddick et al., 2009). The brain DA neurotransmitter system is under a delicate balance between dopamine release, reuptake, and metabolism (Musacchio J. M., 2013). Built upon findings in this study, it is our premise that research that can identify changes in DA release and reuptake in specific brain regions will provide important details about the underlying mechanistic pathways associated with many human diseases. Replication with other mammalian and non-mammalian species with functional monoamine systems in the brain will help assess the reproducibility of these results (Dahlbom et al., 2012; Hall et al., 2004; Kabelik et al., 2014). In this initial study, we focused on the brain dopaminergic system. Future studies using systematic biology approaches will enrich our understanding of the effects of social ranks. Here we show the feasibility of a social rank pig model. In order to provide more definitive evidence for drug use, a drug self-administration component will be necessary.

A major outcome from this study was data showing that the pig is a suitable species to study social enrichment, social stress and the central nervous system function. Social stress is common in humans and has been linked to various mental and behavioral outcomes (Björkqvist, 2001; Mantsch et al., 2016).

Nonetheless, definitive evidence for a causal relationship and its underlying mechanisms is hampered by limitations in human research. For research on the course of alcohol- and drug-related problems, there are additional restrictions about providing psychoactive drugs to drug-naïve people. While pig models have been underutilized in biomedical research (Broom, 2010; Gieling et al., 2011), results from the present study show that the pig model has the potential to fill gaps in our knowledge for the following reasons: 1) pigs are social animals that resemble humans in certain facets of their behavioral repertoire, and there are developmental, anatomic, chemical, and structural similarities between pig and human brains (Broom, 2010; Gieling et al., 2011); and 2) pig models have advantages in cost and accessibility over nonhuman primate models, and have advantages in the ability to study nuances of complex social behaviors over rodent models. In addition, pigs grow into adults at a faster pace than nonhuman primates, which provide a good opportunity to study the effects of childhood or adolescent events on consequences during the adulthood.

The main limitation of the study is missing data due to unsuccessful CSF sample collections. However, it is worth noting that success in obtaining CSF samples was not associated with CSF DA levels or changes in social rank. In this initial study, we only included just-weaned male piglets. Generalization of these findings to females or adults is premature (J. D. Higley et al., 1991).

## 4. Experimental Procedures and Statistical Analysis

### 4.1 Experimental Procedures

The subjects were 16 just-weaned experimentally naïve male piglets (Sus scrofa), 21 days old. Throughout the experiment, all piglets were fed regularly, and water was available ad libitum. Animal housing, handling and all experimental procedures were performed in accordance with the 2011 National Research Council Guidelines for the Care and Use of Mammals in Neuroscience and Behavioral Research and were approved by the Animal Care and Use Committee of Michigan State University.

The experiment utilized a crossover rank-and-re-rank design. During phase I of the experiment (days 1 to 14), 16 just-weaned piglets from different litters were randomly assigned to social groups of four piglets after being stratified by weight at baseline to avoid potential confounding by initial weight (see Figure 1). In each of the four groups, there was one dominant- (the α), one subordinate- (the δ), and two intermediate- (β and γ) ranked piglets. Piglets were housed together for two weeks during Phase I of the experiment (i.e., the initial post-weaning social ranking phase). During Phase II of the experiment (the re-rank phase, days 15 to 28), the four piglets with the same rank from each of the four initial groups (i.e., the four α pigs, β pigs, γ pigs and δ pigs) were placed together on the 15^th^ day of the experiment in order to form new social hierarchies.

**Figure 1.**
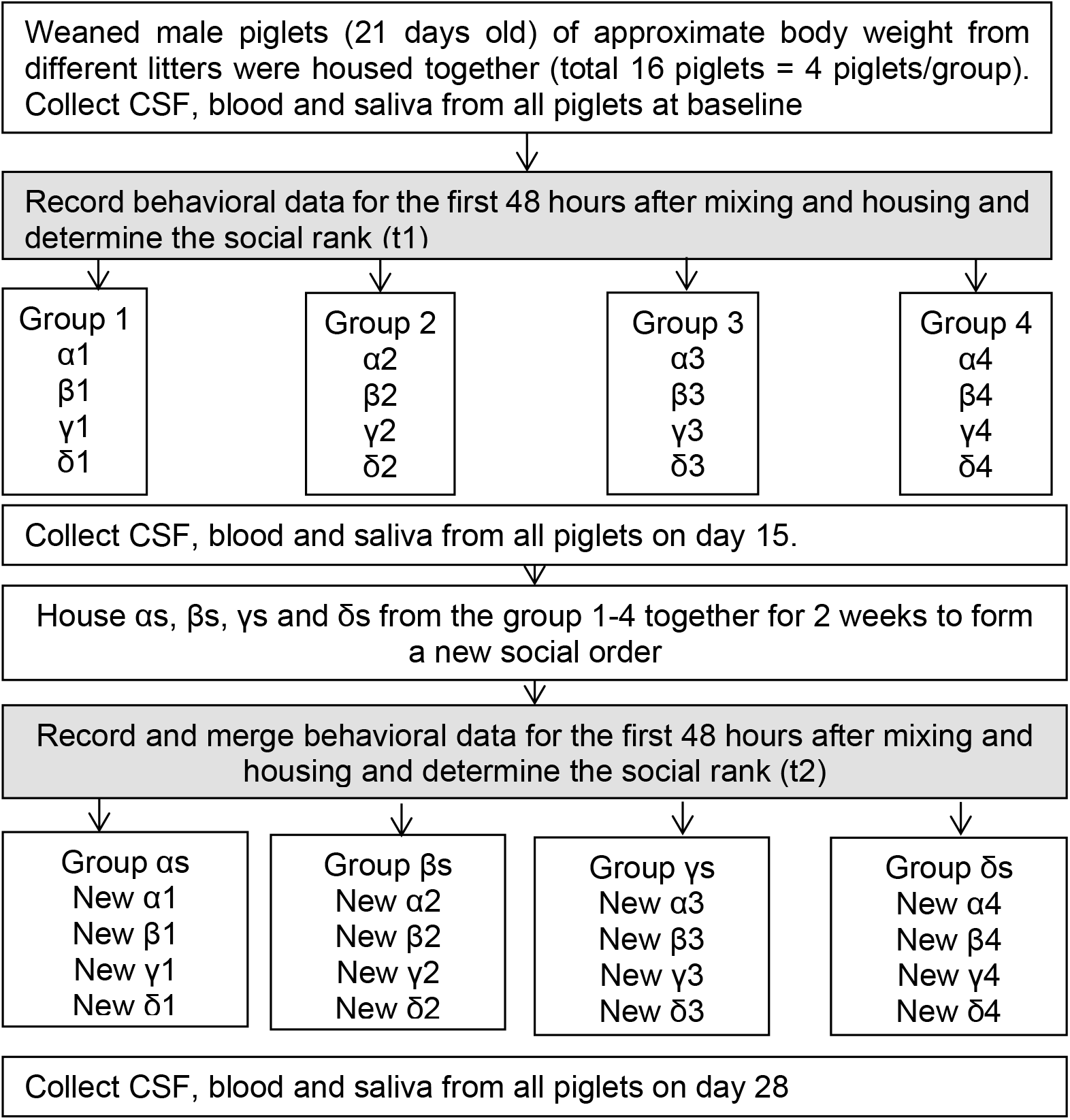
Depiction of study design.

**Figure 2.**
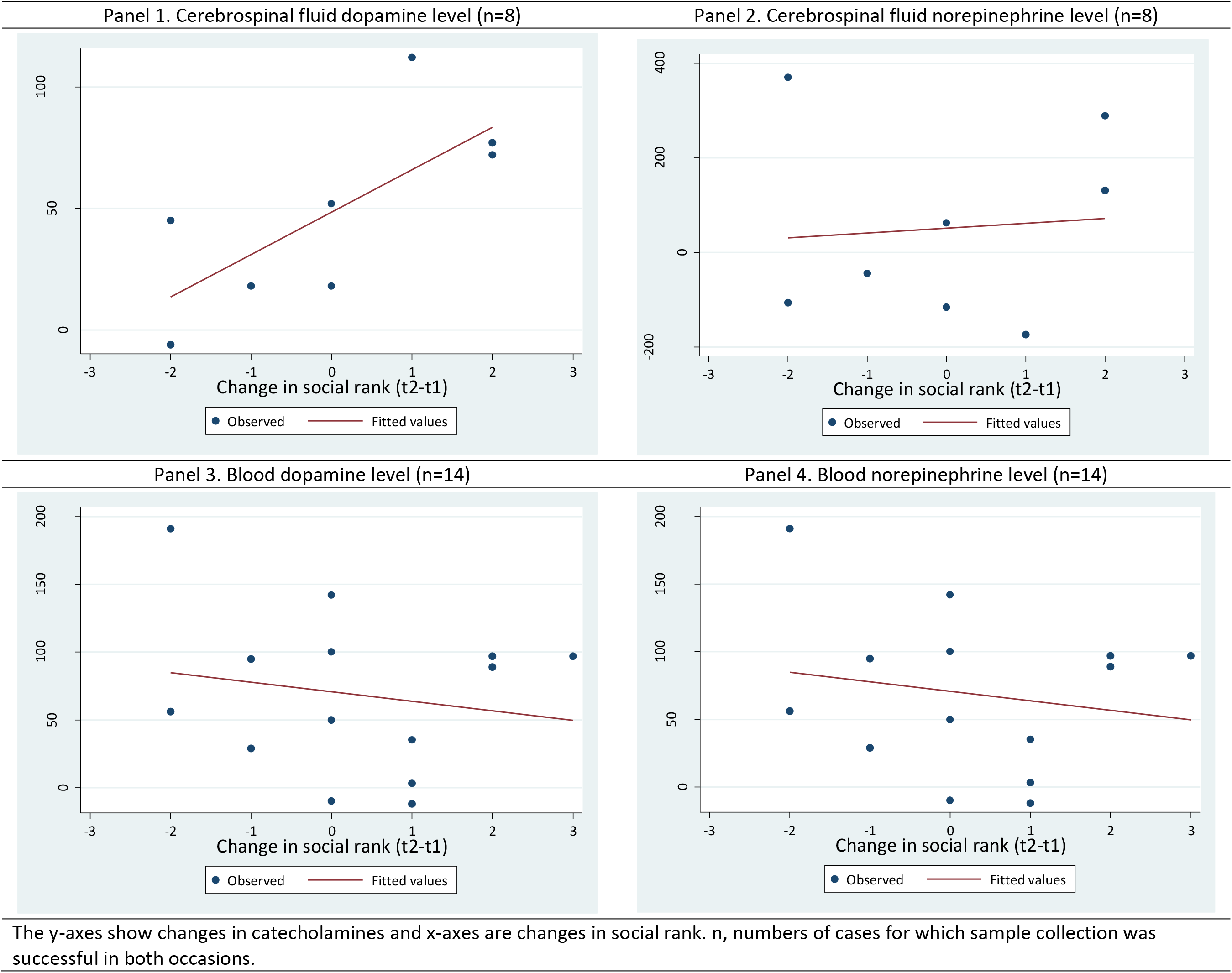
Scatter plots showing the relationship of changes in catecholamines (pg/ml) and changes in social rank in pigs. Panel 1. Cerebrospinal fluid dopamine level (n=8). Panel 2. Cerebrospinal fluid norepinephrine level (n=8). Panel 3. Blood dopamine level (n=14). Panel 4. Blood norepinephrine level (n=14). The y-axes show changes in catecholamines and x-axes are changes in social rank. n, numbers of cases for which sample collection was successful in both occasions.

#### Social Rank Determinations

During the entire experiment, pig behaviors were recorded using an overhead video camera with GeoVision 1480 software. Continuous behavioral sampling was used to collect the following agonistic interaction data during the first 48 hours of each round of group formation (i.e., the first 48 hours during Phase I and Phase II): identification of the piglet who initiated the interaction; the piglets that were involved in the interaction; offensive (e.g., bites, head thrusts, chase, threat, and replacement) and defensive (e.g., freeze, avoidance, and flight) behaviors. To establish social hierarchy, agonistic behaviors occur most frequently during the first 48 hours; once social hierarchy is established, it remains stable for the rest of the study phase. The outcome of the interaction from each pair in the group was calculated as a win or loss. The data were placed in a matrix based on the number of wins a specific piglet had against another piglet. From the matrix, the total wins and losses for each piglet were tabulated and a ratio of wins to losses was calculated for each piglet. The piglet with the highest ratio was assigned the highest ranking (dominant or α), and the piglet with the lowest ratio was assigned the lowest ranking (subordinate or δ) and the remaining two piglets were designated intermediate rankings (β and γ). The same scoring procedure was used in Phase II to determine new social rankings.

#### Biomarker Assessments

Weight, CSF and blood samples were collected from all 16 piglets at the time of weaning (i.e., day one before social housing with piglets from other litters, baseline), at day 15 (i.e., after the first round of ranking and right before the second round of ranking), and day 28 (i.e., after the second round of ranking) of the experiment. In this study, our focus is the effects of social rank instead of the initial agonistic behaviors which are required to the establishment of social rank. Agonistic behaviors usually subside after 48 hours once social ranks are established. We collected CSF and blood samples after two weeks of social housing in order to eliminate any potential confounding effects due to agonistic behaviors and to assess catecholamine concentrations in stable social groups.

Blood samples were collected from the ear vein without sedation using K2-EDTA as an anticoagulant, and the samples were inverted several times to ensure adequate mixing of blood and anticoagulant. After the collection of blood samples, pigs were anesthetized using ketamine (20 mg/kg, xylazine) and midazolam (0.4 mg/kg). Cerebrospinal fluid (CSF) samples were collected from the atlanto-occipital space and stored at −80°C until analyzed. Blood samples were kept on ice during sample collection and centrifuged to harvest plasma afterwards. Plasma was stored at −80°C until analyzed and underwent catecholamine extraction procedure. Briefly, 100 μl of plasma was mixed with 7 mg of activated alumina, 0.4 ml of 2M Tris and 0.5M EDTA PH 8.1, and 15 μl of internal standard (3,4-Dihydroxybenzylamine). The mixture was spun on a vortex for 20 min to attach the catecholamines to the alumina. After washing the alumina, catecholamines were eluted from the alumina with 100 μl of 0.2M Acetic Acid. The concentration of DA, NE and epinephrine were determined by high performance liquid chromatography with electrochemical detection and expressed as pg/ml.

CSF samples were successfully collected from eight pigs at both time points after the first and second social ranking, and blood samples were successfully collected from 15 pigs at both time points. Whether CSF sample collection was successful or not was not associated with CSF dopamine or norepinephrine levels at all three time points or changes in social ranks (odds ratio=1.0 for all measures; all p>0.35). One pig was excluded from the main analysis due to extreme values in blood catecholamine levels. CSF samples were not collected after the second round of ranking for this pig and therefore, it was not included in the CSF analysis.

### 4.2 Statistical analysis

First, we tested the normality of data using a joint test for skewness and kurtosis (Royston, 1991). CSF epinephrine concentrations were too low to be detected by high performance liquid chromatography, and therefore could not be analyzed. Linear regression was used to estimate whether baseline weight and catecholamine levels predicted the first social rank. A dominant rank was coded “4”, and a subordinate rank was coded “1” with intermediate ranks coded “2” and “3”. The change in social rank between the two rounds of ranking was calculated by subtracting the rank during the first ranking from the rank during the second ranking. Therefore, a positive number indicates an increase in social, a negative number represents a drop in social rank, and “0” indicates no change in social rank. Similarly, changes in catecholamine levels were calculated by subtracting the catecholamine measures after the first ranking from the measures after the second ranking. Scatterplots were used to graph changes in catecholamine levels against changes in social rank. Linear regression was used to estimate post-rank changes in catecholamine levels associated with changes in social ranks. In order to explore potential moderation by initial social rank, we categorized piglets into four groups: low rank becoming high rank (n=3 for CSF measures, n=4 for blood measures); low rank remaining low rank (n=2 for CSF measures, n=4 for blood measures), high rank becoming low rank (n=2 for CSF measures, n=4 for blood measures), and high rank remaining high rank (n=1 for CSF measures, n=3 for blood measures). In this exploratory analysis, αs and βs were treated as high ranks; δs and γs are treated as low ranks. Linear regression was used to estimate differences in post-rank changes in catecholamine levels across these groups.

## Acknowledgment

We wish to thank the van der Staay & Nordquist swine research lab at University of Utrecht, Dr. Gregory Fink, Dr. Keith Lookingland, and Dr. David Todem at Michigan State University for providing expert consultation and technical support.

## References

Björkqvist, K., 2001. Social defeat as a stressor in humans. Physiol. Behav. 73, 435–442. doi:10.1016/S0031-9384(01)00490-5

Bowling, S.L., Rowlett, J.K., Bardo, M.T., 1993. The effect of environmental enrichment on amphetamine-stimulated locomotor activity, dopamine synthesis and dopamine release. Neuropharmacology 32, 885–893. doi:10.1016/0028-3908(93)90144-R

Broom, D.M., 2010. Cognitive ability and awareness in domestic animals and decisions about obligations to animals. Appl. Anim. Behav. Sci. 126, 1–11. doi:10.1016/j.applanim.2010.05.001

Clarke, A.S., Hedeker, D.R., Ebert, M.H., Schmidt, D.E., McKinney, W.T., Kraemer, G.W., 1996. Rearing experience and biogenic amine activity in infant rhesus monkeys. Biol. Psychiatry 40, 338–352. doi:10.1016/0006-3223(95)00663-X

Crowley, T.J., Mikulich, S.K., Williams, E.A., Zerbe, G.O., Ingersoll, N.C., 1992. Cocaine, social behavior, and alcohol-solution drinking in monkeys. Drug Alcohol Depend. 29, 205–223. doi:10.1016/0376-8716(92)90094-S

Crowley, T.J., Stynes, A.J., Hydinger, M., Kaufman, I.C., 1974. Ethanol, methamphetamine, pentobarbital, morphine, and monkey social behavior. Arch. Gen. Psychiatry 31, 829–838. doi:10.1001/archpsyc.1974.01760180069009

Czoty, P.W., Gould, R.R., Gage, H.D., Nader, M.A.. n.d., Effects of Social Reorganization on Dopamine D2/D3 Receptor Availability and Cocaine Self-Administration in Male Cynomolgus Monkeys. under Rev.

Czoty, P.W., McCabe, C., Nader, M. a, 2005. Assessment of the relative reinforcing strength of cocaine in socially housed monkeys using a choice procedure. J Pharmacol Exp Ther 312, 96–102. doi:10.1124/jpet.104.073411

Czoty, P.W., Morgan, D., Shannon, E.E., Gage, H.D., Nader, M. a, 2004. Characterization of dopamine D1 and D2 receptor function in socially housed cynomolgus monkeys self-administering cocaine. Psychopharmacology (Berl). 174, 381–8. doi:10.1007/s00213-003-1752-z

Dahlbom, S.J., Backström, T., Lundstedt-Enkel, K., Winberg, S., 2012. Aggression and monoamines: Effects of sex and social rank in zebrafish (Danio rerio). Behav. Brain Res. 228, 333–338. doi:10.1016/j.bbr.2011.12.011

Dohrenwend, B.P., Levav, I., Shrout, P.E., Schwartz, S., Naveh, G., Link, B.G., Skodol, a E., Stueve, A., 1992. Socioeconomic status and psychiatric disorders: the causation-selection issue. Science (80-.). 255, 946–952. doi:10.1126/science.1546291

Gieling, E.T., Schuurman, T., Nordquist, R.E., van der Staay, F.J., 2011. The pig as a model animal for studying cognition and neurobehavioral disorders, in: Hagan, J.J. (Ed.), Current Topics in Behavioral Neurosciences. Global Medical Excellence Cluster (GMEC), United Kingdom, pp. 359–383. doi:10.1007/7854_2010_112

Gill, K.E., Pierre, P.J., Daunais, J., Bennett, A.J., Martelle, S., Gage, H.D., Swanson, J.M., Nader, M. a, Porrino, L.J., 2012. Chronic Treatment with Extended Release Methylphenidate Does Not Alter Dopamine Systems or Increase Vulnerability for Cocaine Self-Administration: A Study in Nonhuman Primates. Neuropsychopharmacology 37, 2555–2565. doi:10.1038/npp.2012.117

Greene, K.A., Faull, K.F., 1989. Relationship between plasma and cerebrospinal fluid norepinephrine and dopamine metabolites in a nonhuman primate. J Neurochem 53, 1007–1013.

Hall, F.S., Sora, I., Drgonova, J., Li, X.F., Goeb, M., Uhl, G.R., 2004. Molecular mechanisms underlying the rewarding effects of cocaine, in: Annals of the New York Academy of Sciences. doi:10.1196/annals.1316.006

Harlow, H.F., Zimmermann, R.R., 1959. Affectional responses in the infant monkey; orphaned baby monkeys develop a strong and persistent attachment to inanimate surrogate mothers. Science (80-.). 130, 421–432. doi:10.1126/science.130.3373.421

Hemmingsson, T., Lundberg, I., Diderichsen, F., 1999. The roles of social class of origin, achieved social class and intergenerational social mobility in explaining social-class inequalities in alcoholism among young men. Soc. Sci. Med. 49, 1051–1059. doi:10.1016/S0277-9536(99)00191-4

Higley, J D, Hasert, M.F., Suomi, S.J., Linnoila, M., 1991. Nonhuman primate model of alcohol abuse: effects of early experience, personality, and stress on alcohol consumption. Proc. Natl. Acad. Sci. U. S. A. 88, 7261–7265. doi:10.2307/2357662

Higley, J. D., Suomi, S.J., Linnoila, M., 1991. CSF monoamine metabolite concentrations vary according to age, rearing, and sex, and are influenced by the stressor of social separation in rhesus monkeys. Psychopharmacology (Berl). doi:10.1007/BF02244258

Kabelik, D., Alix, V.C., Singh, L.J., Johnson, A.L., Choudhury, S.C., Elbaum, C.C., Scott, M.R., 2014. Neural activity in catecholaminergic populations following sexual and aggressive interactions in the brown anole, Anolis sagrei. Brain Res. 1553, 41–58. doi:10.1016/j.brainres.2014.01.026

Kanitz, E., Puppe, B., Tuchscherer, M., Heberer, M., Viergutz, T., Tuchscherer, A., 2009. A single exposure to social isolation in domestic piglets activates behavioural arousal, neuroendocrine stress hormones, and stress-related gene expression in the brain. Physiol. Behav. 98, 176–185. doi:10.1016/j.physbeh.2009.05.007

Kanitz, E., Tuchscherer, M., Puppe, B., Tuchscherer, A., Stabenow, B., 2004. Consequences of repeated early isolation in domestic piglets (Sus scrofa) on their behavioural, neuroendocrine, and immunological responses. Brain. Behav. Immun. 18, 35–45. doi:10.1016/S0889-1591(03)00085-0

Koob, G.F., Le Moal, M., 1997. Drug abuse: hedonic homeostatic dysregulation. Science (80-.). 278, 52–58.

Koob, G.F., Volkow, N.D., 2010. Neurocircuitry of addiction. Neuropsychopharmacology 35, 217–238. doi:10.1038/npp.2010.4

Link, B.G., Phelan, J., 1995. Social Conditions As Fundamental Causes of Disease Social Conditions as Fundamental Causes of Disease*. Source J. Heal. Soc. Behav. J. Heal. Soc. Behav. 80–9480. doi:10.2307/2626958

Mantsch, J.R., Baker, D.A., Funk, D., Lê, A.D., Shaham, Y., 2016. Stress-induced reinstatement of drug seeking: 20 years of progress. Neuropsychopharmacology. doi:10.1038/npp.2015.142

Marmot, M., 2005. Social determinants of health inequalities. Lancet 365, 1099–1104. doi:10.1016/S0140-6736(05)71146-6

Morgan, D., Grant, K. a, Prioleau, O. a, Nader, S.H., Kaplan, J.R., Nader, M. a, 2000. Predictors of social status in cynomolgus monkeys (Macaca fascicularis) after group formation. Am. J. Primatol. 52, 115–131. doi:10.1002/1098-2345(200011)52:3<115::AID-AJP1>3.0.CO;2-Z</115::AID-AJP1>

Morgan, D., Grant, K.A., Gage, H.D., Mach, R.H., Kaplan, J.R., Prioleau, O., Nader, S.H., Buchheimer, N., Ehrenkaufer, R.L., Nader, M.A., 2002. Social dominance in monkeys: dopamine D2 receptors and cocaine self-administration. Nat. Neurosci. 5, 169–74. doi:10.1038/nn798

Musacchio J. M., 2013. Enzymes involved in the biosynthesis and degradation of catecholamines, in: L., I. (Ed.), Biochemistry of Biogenic Amines. Springer., pp. 1–35.

Nader, M A, Czoty, P.W., Nader, S.H., Morgan, D., 2012. Nonhuman primate models of social behavior and cocaine abuse. Psychopharmacol. 224, 57–67. doi:10.1007/s00213-012-2843-5

Nader, Michael A., Nader, S.H., Czoty, P.W., Riddick, N. V., Gage, H.D., Gould, R.W., Blaylock, B.L., Kaplan, J.R., Garg, P.K., Davies, H.M.L., Morton, D., Garg, S., Reboussin, B.A., 2012. Social dominance in female monkeys: Dopamine receptor function and cocaine reinforcement. Biol. Psychiatry 72, 414–421. doi:10.1016/j.biopsych.2012.03.002

Nader, M.A., Woolverton, W.L., 1991. Effects of increasing the magnitude of an alternative reinforcer on drug choice in a discrete-trials choice procedure. Psychopharmacology (Berl). 105, 169–74. doi:10.1007/BF02244304

Nobile, M., Giorda, R., Marino, C., Carlet, O., Pastore, V., Vanzin, L., Bellina, M., Molteni, M., Battaglia, M., 2007. Socioeconomic status mediates the genetic contribution of the dopamine receptor D4 and serotonin transporter linked promoter region repeat polymorphisms to externalization in preadolescence. Dev. Psychopathol. 19, 1147–1160. doi:10.1017/S0954579407000594

Riddick, N. V., Czoty, P.W., Gage, H.D., Kaplan, J.R., Nader, S.H., Icenhower, M., Pierre, P.J., Bennett, A., Garg, P.K., Garg, S., Nader, M.A., 2009. Behavioral and neurobiological characteristics influencing social hierarchy formation in female cynomolgus monkeys. Neuroscience 158, 1257–1265. doi:10.1016/j.neuroscience.2008.11.016

Royston, P., 1991. Comment on sg3.4 and an improved D’Agostino test. Stata Technical Bulletin 3: 23-24. Stata Tech. Bull. Repr. 1, 110–112.

Tung, J., Akinyi, M.Y., Mutura, S., Altmann, J., Wray, G.A., Alberts, S.C., 2011. Allele-specific gene expression in a wild nonhuman primate population. Mol. Ecol. 20, 725–739. doi:10.1111/j.1365-294X.2010.04970.x

Volkow, N.D., Fowler, J.S., Wang, G.-J., 1999. Imaging studies on the role of dopamine in cocaine reinforcement and addiction in humans. J. Psychopharmacol. 13, 337–345. doi:10.1177/026988119901300406

Volkow, N.D., Morales, M., 2015. The Brain on Drugs: From Reward to Addiction. Cell 162, 712–725. doi:10.1016/j.cell.2015.07.046

